# Using a Champion-Oriented Mindset to Overcome the Challenges of Graduate School

**DOI:** 10.1101/2021.11.29.469904

**Authors:** Andrea G. Marshall, Caroline B. Palavicino-Maggio, Kit Neikirk, Zer Vue, Heather Beasley, Edgar Garza-Lopez, Sandra Murray, Denise Martinez, Jamaine Davis, Haysetta Shuler, Elsie C. Spencer, Derrick Morton, Antentor Hinton

**Author notes:** These authors share co-first authorship (Andrea G. Marshall, Caroline B. Palavicino-Maggio). These authors share co-senior authorship (Haysetta Shuler, Elsie C. Spencer, Derrick Morton, Antentor Hinton).

## Abstract

Despite efforts to increase diversity, a glaring underrepresentation of minorities (URM) persists in the fields of science, technology, engineering, and mathematics (STEM). Graduate school can be a stressful step in the STEM pipeline, especially for students previously unaware of the structure and challenges of post-graduate education. To promote successful minority participation in STEM and prepare prospective students for the impending challenges of graduate school, we developed a workshop based on the mentoring and fostering of a champion-oriented mindset entitled, “The Trials and Tribulations of Graduate School: How Do You Make an Impact?”. We administered the workshop to a cohort of university undergraduates and conducted pre- and post-workshop surveys to measure students’ perceived need for instruction on specific workshop topics. The results suggest that the workshop was well received by the students and provided information that they considered helpful to help navigate the graduate school process.

## Introduction

Despite many institutional initiatives and programs aimed to increase the presence of underrepresented minorities (URMs) in science, technology, engineering, and mathematics (STEM), URM participation in STEM fields still remains disproportionately low, in part because support programs often have limited availability and therefore fail to reach these pool of students [1, 2]. Here, we define URMs as Black or African Americans, Latinx or Hispanic Americans, American Indians, Alaskan Natives, Native Hawaiians, and Native Pacific Islanders [3]. According to the National Science Foundation (NSF) and the National Science Board (NSB), there is a critical need to bolster the presence and retention of URMs in STEM. Diversity in STEM is important because multiformity leads to more impactful research, improved problem solving skills, decreased turnover rates [4], and promotes elevated innovation [3]. Moreover, the number of STEM occupations is expected to increase at almost twice the rate of overall job growth in the U.S. [1, 2]. This can potentially lead to even greater racial disparities unless intentional efforts are increased to promote and sustain URM participation in STEM. As part of these efforts, we believe that faculty should take an active role by providing mentorship, support, and advocacy for URM students pursuing STEM careers.

Graduate- and post-graduate level education is required for many STEM occupations. Beyond the daily stresses of graduate school, URMs face additional challenges that can hinder their productivity and deter them from pursuing or obtaining a career in STEM, including microaggressions [5], imposter syndrome [3, 5, 6], awfulizing and demoralization [7], and lack of or decreased access to mentors [4]. Importantly, affirmative mentoring [8-10], access to career development resources, and self-efficacy [11-13] are correlated with positive academic and professional outcomes in STEM [14, 15] [16]. Some organizations including the National Institutes of Health (NIH) have programs like the Maximizing Access to Research Careers (MARC) and Research Initiative for Scientific Enhancement (RISE) training grants, that seek to enhance and promote URM presence in STEM by providing financial support as well as research, training, professional development, and mentoring activities to help students prepare for graduate school (https://www.nigms.nih.gov/training/marc/pages/ustarawards.aspx). However, these programs are not available at every university and the number of opportunities available for each training grant is limited, thus curtailing the overall reach of their impact. To help close this gap and promote graduate school preparedness among as many URM students in STEM as possible, we developed a workshop based on the framework of intentional mentoring. Participants in the workshop are trained how to increase their self-efficacy skills by adopting a “champion’s mindset” to overcome the hurdles that they will face in the STEM pipeline.

### Framework for the Workshop

Here, we present the content of the workshop regarding what undergraduate students should expect in graduate school and how they can maximize their chances of success in a graduate program. Students are surveyed before the workshop and after its completion.

### The Champion’s Mindset

In the workshop, participants are presented a model in which intelligence is viewed as either static and unchangeable (fixed mindset) or flexible with room for development (champion’s mindset; **Figure 1**)[17, 18]. Individuals with a fixed mindset tend to be fragile and avoid challenges because they view them as threats that might reveal inherent personal deficiencies. These individuals also interpret expenditure of effort as a sign of low capacity and setbacks as evidence of low ability [19]. By contrast, individuals with a champion’s mindset innately seek out challenges because they perceive themselves as able to conquer them, create opportunities for improvement, and better hone their abilities [19]. Research has shown that students at all levels who approach learning with a champion’s mindset learn more, achieve more, and perform better academically than students who approach learning with a static mindset [19]. Therefore, our workshop emphasizes the importance of adopting a champion’s mindset as they enter a graduate program.

**Figure 1.**
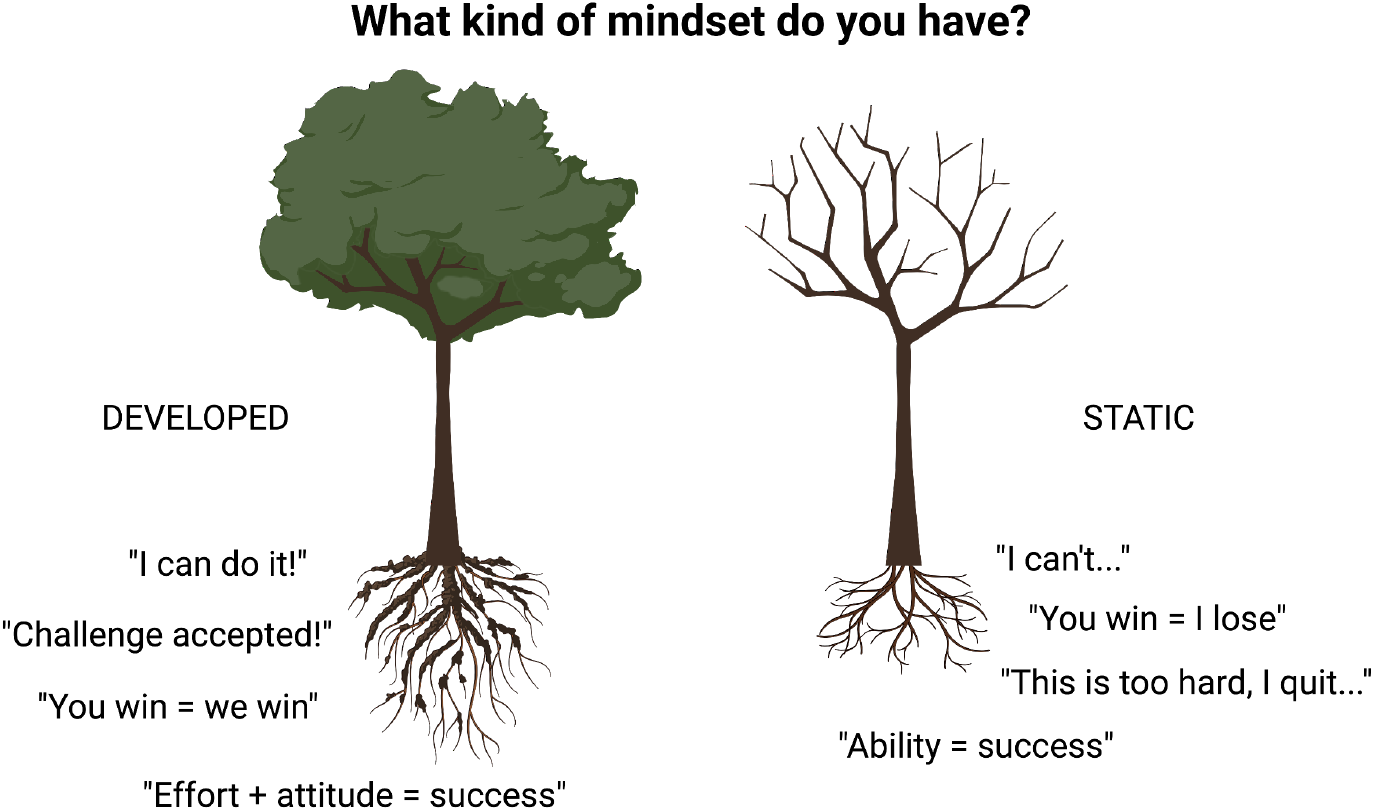
Infographics from the workshop depicting growth or developed mindsets compared to fixed or static mindsets. Someone with a growth mindset tries to work with others, as they understand that success in STEM is not dependent on one person, whereas those with a fixed mindset can avoid collaborating because they see another person succeeding as themselves losing.

### Early Exposure to Undergraduate Research Experiences

Early exposure to STEM experiences can significantly influence a student’s motivation and preparedness to pursue a career in STEM [3, 4, 20]. Therefore, students who participated in the workshop were encouraged to seek opportunities to conduct research prior to applying or attending graduate school through programs such as the NIH’s T34 Bridges to the Baccalaureate Research Training Program, Summer Research Opportunity Program, or Research Experiences for Undergraduates [3]. The workshop emphasized that research experiences can occur both within and outside the student’s home institution. However, it is important to underscore that because many minority-serving institutions (MSI’s) lack the research infrastructure and funds to support such programs, it is critical that students be made aware of the existence of these programs at other schools.

### Mentors Are a Necessity

Another key element highlighted in the workshop is the power and importance of mentors. Effective mentoring is personal and reciprocal in nature. It helps the mentee achieve their goals, and provides psychosocial, professional, and career support [7]. Importantly, mentoring can mitigate the pitfalls that URMs in graduate programs commonly face, including burnout, minority stress, imposter syndrome, demoralization, and awfulization [3, 5-7]. This support is accomplished through intentionality and purpose: the mentors and mentees are deliberate in getting to know one another, communicate openly and frequently, continually reflect on the mentor-mentee relationship, and work in unison to achieve the mentee’s goals. This is implemented by utilizing tools such as mentoring contracts and individual development plans (IDP’s) [7]. In our workshop, students are provided with practical advice on choosing mentors who are genuinely committed to their success, invested in their long-term development, and able to focus on their strengths while addressing their weaknesses in a proactive manner. Students are also advised to seek multiple mentors with different strengths, rather than limiting themselves to only one mentor. We understand that the process of identifying effective mentors can take time, but we strongly encourage students to look for role models as well, such as people outside of the institution who are successful practitioners in an area (or areas) of interest to them and/or those who have overcome adversity themselves. Even when students lack a mentor, they can still use proactive (rather than reactive) development options to effectively prepare for graduate school. For example, students can read about their role model’s journey and learn what steps that person took to become successful including past grants they were awarded and various public or community engagements they were involved in during their careers. This will allow a mentee to gauge how well that individual lead, inspired, and cultivated other future URMs in STEM fields.

### Distinguishing Oneself in Graduate School Applications

Specific application requirements vary by university and program. Typically applications require official transcripts detailing the applicant’s overall grade point average (GPA) and possibly the GPA within the subject area of the applicant’s major; class ranking; a condensed personal statement; graduate record examination (GRE) scores (although some programs waive this requirement); multiple letters of recommendation; and a resume or curriculum vitae (CV). Understanding the relevance a CV plays into the application process is critical. The CV should indicate all honors and awards received by the applicant (e.g., fellowships, scholarships, travel awards), relevant job experience (e.g., internships, research, teaching), participation in campus organizations (e.g., society memberships, sports activities, club memberships), community service, scholarship, and leadership activities. The minimum GPA requirement for many Ph.D. programs is 3.0 or equivalent to a B-. If their GPA is lower than 3.0, the applicant may consider retaking courses (if applicable), applying to a post-baccalaureate program, or pursue a master’s degree before applying to a Ph.D. program. This suggestion will not be frowned upon by admissions committees but will highlight the willingness and determination that an applicant has in pursuing a doctoral degree in STEM. GRE scores within the 75^th^ percentile are acceptable for most schools; however, scores in the 90^th^ percentile or above are desirable and typically accepted by all schools.

The personal statement should be the applicant’s “story” or positionality that draws in the reader and motivates them to meet the applicant in person. Therefore, word choice is critical. Effective personal statements usually include an exciting hook or angle that distinguishes the author from other applicants. It explains why the individual wants to pursue a Ph.D. degree in general and what they intend to do with that degree from that institution should they be accepted. In general, all applicants to competitive schools have exceptional GPAs and GRE scores, so the personal statement should include information that makes the candidate stand out among other qualified individuals. Things that stand out are involvement in extracurricular activities, volunteer or community work, and publications. The personal statement should also explain why the applicant chose to apply to the specific school and program. This is an opportunity for admissions committee members to see how much you understand the inner workings of their institutions and how well aligned you are to their mission. Furthermore, it is acceptable to discuss both strengths and weaknesses in the personal statement, however, the applicant should be careful to avoid coming across as overconfident or unqualified.

Students should be forward-thinking when identifying individuals from whom they may ask for a letter of recommendation. Someone who knows the applicant well (e.g., their strengths and weaknesses, struggles and triumphs, and work ethic), can likely write a more personalized and ultimately effective letter. Hence, it is generally not recommended for students to ask for a letter of recommendation from a professor who they have only seen in a classroom filled with hundreds of other students. The best way to obtain effective letters of recommendation is to establish and maintain good relationships with professors, mentors, and research associates before applying to graduate programs. In this scenario, when applying to graduate programs, students can more readily request letters from people who know them well and can speak to specific qualities that make them an ideal candidate. It is important to emphasize that time is of critical essence when requesting a letter of recommendation. An applicant typically should give the author a minimum of 4–6-weeks’ notice before the submission deadline when requesting a letter of recommendation. Phrasing is key when asking a potential recommender. It should not be assumed that a recommender will be willing to provide one or able to within the allotted time. But if they agree, which they normally do, students should for a strongly written, *positive* letter of recommendation addressed to their program of choice. The student should then pay close attention to the nuances of the answer because a recommendation can be positive, negative, or neutral. Once a person agrees to write a strong, affirming letter, the student should provide them with a copy of transcripts, a reminder of pertinent shared experiences, an outline of their goals, and a CV detailing all relevant research experience, internships, leadership activities, awards, and achievements that can be referenced in the letter as needed.

Before applying to graduate school, and after the application process, it is important for students to reflect on what might be beneficial for their mental health. This is a critical step that often gets overlooked prior and during the application process. For example, students may find it useful to explore their personality type or identify activities that restore their energy by using tools such as the Big Five Personality Test, the Myers-Briggs Type Indicator, or the Five Love Languages. Students should expect graduate school to be challenging, not just intellectually but in all aspects of their life. Hence, the better they know themselves and their unique learning styles, the more prepared they will be to overcome and address the challenges of pursuing a graduate education head-on.

### Interviewing Prospective Graduate Programs

Applicants should remember that they are assessing graduate programs just as much as graduate programs are assessing them. They should therefore proactively research the institutions that will be their home away from home for the next five to six years. This research should also include academic departments, the clinical affiliations of the institution (if any), and the cities and neighborhoods surrounding the institution. Some questions to consider during this process include: What types of communities exist that might be a good fit for the applicant? Are current URM graduate students happy or satisfied with the program and school? Are resources available to accommodate and support the specific needs of the applicant? What is the faculty to student ratio? These questions are important to consider because graduate school can be a prolonged season of both personal and professional trials, therefore, applicants should use their interviews with prospective graduate programs as opportunities to gauge whether each program is right for them. They should truly ask themselves whether that institution and its culture and environment could be a place in which the individual may develop a sense of community, which will in turn help them to succeed.

The interview phase is another process of the graduate school process that students need to be prepared to successfully undergo. Regardless of whether the interview is on-site or virtual, the applicant must be enthusiastic and convey curiosity, competence, and courtesy. Unless otherwise stated, the expected attire for in-person interviews is generally semi-formal but comfortable enough to endure the several hours required to complete the interview process, which is also referred to as “running the gauntlet”. Before the day of the interview, applicants should familiarize themselves with the faculty in the department, their research interests and accolades, and how the applicant will fit into that environment. Applicants should also identify three to five potential mentors at that institution who they would like to meet during their visit and potentially work with for the duration of graduate school.

### The Structure of Graduate School

Before starting graduate school, students should rest, recharge, refresh, and engage in a champion’s mindset. **Table 1** shows the general structure and suggested goals for each year of graduate school. Typically, the first two years are the most challenging, during which students usually complete most of their required academic coursework (typically at a faster pace than undergraduate courses). They rotate through laboratories, often shadowing senior researchers, principal investigators, and faculty and select one in which to complete their graduate research, and complete their qualifying exam. Throughout these early years, students should be proactive in getting to know their professors and working together with their cohort to excel in courses. Because balancing coursework, relationship building, and laboratory rotations is difficult, it is crucial to engage in self-reflection and stress management activities at this stage of their academic career. Moreover, it is not uncommon for relationship dynamics with family and friends to change during these years, as graduate school is often more taxing than undergraduate studies. Students and their families have to understand that social events and familial commitments will need to be adjusted due to the student’s schedule and workload. For many students, the first two years of graduate school are a transition period during which they shift from mostly following instructions to making their own informed decisions.

**Table 1.** Overview and general structure of graduate school.

| Year | Focus |
| --- | --- |
| 1 | General Program Coursework and Laboratory Rotations |
| 2 | Specialized Coursework and Qualifying Exam |
| 3 | Career Development and Grant Writing |
| 4 | Collaboration and Publications |
| 5+ | Next Career Steps and Graduation |

Another challenge in the graduate school phase is the qualifying exam (QE), an oral exam that determines whether or not students will be allowed to continue in their graduate program after completion of their first two years. This exam is a requirement for all Ph.D. candidates and assesses whether a student is academically prepared for the following years of instruction. Generally, this exam involves an examination committee comprised of three faculty who will ask questions on your proposed research topic then make suggestions and recommendations to strengthen or clarify your research focus. In preparation for this exam, students should be diligent in determining what the exam requires, its format, and schedule sufficient time to study. The conclusion of the QE will result in a committee’s suggestion of pass or fail. Most programs have strict rules for what constitutes a pass, conditional pass, or failure, and how many times the exam may be re-taken before a student who fails to pass is dismissed from the program. Students should prioritize QE preparation and allow for ample study time. They should also become familiar with the specific QE rules and content according to their particular program and school by attending similar venues such as doctoral defenses, which are generally open to the public.

After successfully passing the QE, the student begins year three of the graduate program. During this year students should consider immersing themselves in career development and grant writing opportunities at their home institution and at subject-relevant organizations. During year four, student should focus on writing research manuscripts and establishing networking collaborations, which is critical because many programs require at least one first-author publication for graduation. Throughout and beyond year five, students should consider their next career steps. With guidance from their mentors, they should apply for their next position at least nine to 12 months in advance of when they expect to graduate. This will allow them sufficient time to find a suitable position that aligns with their career goals and to learn any additional skills or experience that may be required for the position. Throughout these years, a student should be expanding their dissertation, which is based on the research topic they used during their QE. Writing each chapter can take months, even years depending on the topic and encapsulates a format that is informative to the reader. It should include research questions, methodology, research framework, discussion, findings, and a conclusion that answers your research questions based on your findings and analyses. In parallel, the process of writing and defending a dissertation or thesis can be daunting, so students should also allow time to correctly format and complete their dissertation by the deadlines set by their school.

## Methods

We administered our 90-minute virtual workshop entitled “The Trials and Tribulations of Graduate School: How to Make an Impact” to 24 undergraduate students at Winston-Salem State University (a historically Black public university) from the general student population. This study was submitted to the Kaiser Permanente review board under the title “Promoting Engagement in science for underrepresented Ethnic and Racial minorities (P.E.E.R),” and was approved and classified as not human subjects research. The participants completed questionnaires before and after the workshop **(see Table 2)** to gauge their expectations and satisfaction regarding the workshop. These were either anonymous paper surveys or an emailed electronic survey. All respondents were informed of the purpose of these surveys, and surveys were on a voluntary basis and independently performed to avoid peer pressure. Respondents that consented to the research returned surveys discreetly in an envelope and/or in an anonymous online form. We summarized the data from the questionnaires using box and whisker plots in which the red centerline denotes the median, and error bars denote the standard error. Individual values are represented by circles. We analyzed the raw data with nonparametric tests for comparison within paired samples. We also used Wilcoxon matched-pairs and signed-rank tests to determine differences between measures. The p-values less than 0.05 indicated statistical significance and NS were non-significant.

**Table 2.**
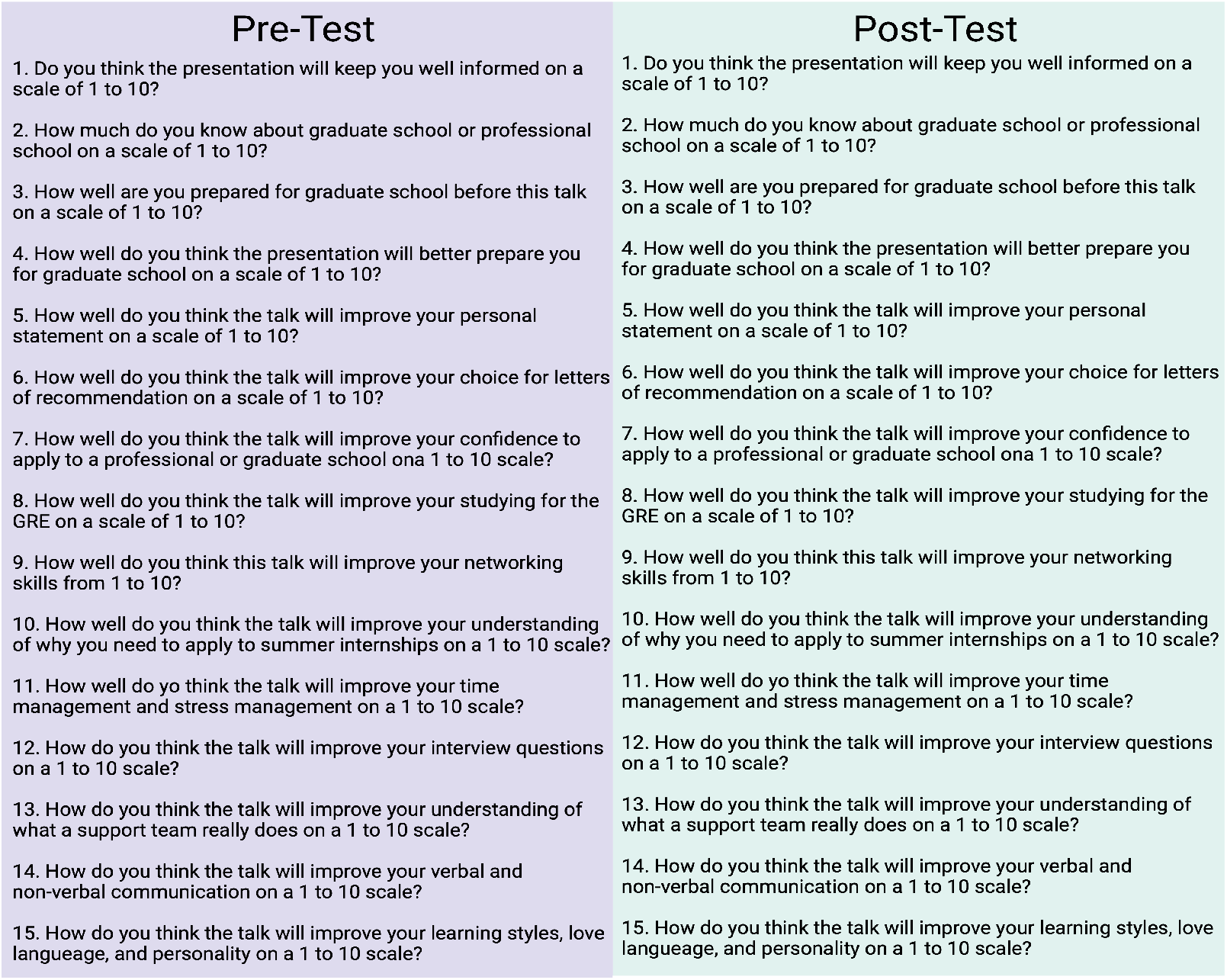
Pre- and post- workshop evaluations.

| Pre-Test | Post-Test |
| --- | --- |
| 1. Do you think the presentation will keep you well informed on a scale of 1 to 10? | 1. Do you think the presentation will keep you well informed on a scale of 1 to 10? |
| 2. How much do you know about graduate school or professional school on a scale of 1 to 10? | 2. How much do you know about graduate school or professional school on a scale of 1 to 10? |
| 3. How well are you prepared for graduate school before this talk on a scale of 1 to 10? | 3. How well are you prepared for graduate school before this talk on a scale of 1 to 10? |
| 4. How well do you think the presentation will better prepare you for graduate school on a scale of 1 to 10? | 4. How well do you think the presentation will better prepare you for graduate school on a scale of 1 to 10? |
| 5. How well do you think the talk will improve your personal statement on a scale of 1 to 10? | 5. How well do you think the talk will improve your personal statement on a scale of 1 to 10? |
| 6. How well do you think the talk will improve your choice for letters of recommendation on a scale of 1 to 10? | 6. How well do you think the talk will improve your choice for letters of recommendation on a scale of 1 to 10? |
| 7. How well do you think the talk will improve your confidence to apply to a professional or graduate school ona 1 to 10 scale? | 7. How well do you think the talk will improve your confidence to apply to a professional or graduate school ona 1 to 10 scale? |
| 8. How well do you think the talk will improve your studying for the GRE on a scale of 1 to 10? | 8. How well do you think the talk will improve your studying for the GRE on a scale of 1 to 10? |
| 9. How well do you think this talk will improve your networking skills from 1 to 10? | 9. How well do you think this talk will improve your networking skills from 1 to 10? |
| 10. How well do you think the talk will improve your understanding of why you need to apply to summer internships on a 1 to 10 scale? | 10. How well do you think the talk will improve your understanding of why you need to apply to summer internships on a 1 to 10 scale? |
| 11. How well do yo think the talk will improve your time management and stress management on a 1 to 10 scale? | 11. How well do yo think the talk will improve your time management and stress management on a 1 to 10 scale? |
| 12. How do you think the talk will improve your interview questions on a 1 to 10 scale? | 12. How do you think the talk will improve your interview questions on a 1 to 10 scale? |
| 13. How do you think the talk will improve your understanding of what a support team really does on a 1 to 10 scale? | 13. How do you think the talk will improve your understanding of what a support team really does on a 1 to 10 scale? |
| 14. How do you think the talk will improve your verbal and non-verbal communication on a 1 to 10 scale? | 14. How do you think the talk will improve your verbal and non-verbal communication on a 1 to 10 scale? |
| 15. How do you think the talk will improve your learning styles, love language, and personality on a 1 to 10 scale? | 15. How do you think the talk will improve your learning styles, love language, and personality on a 1 to 10 scale? |

## Results

### Student Responses and Observations

Responses to the pre-workshop questionnaires suggested that students initially did not see much value in the workshop (**Figure 2, Figure 6**, and **Figure 7**, pre-test). Whether their low expectations were due to a lack of exposure or a lack of mentorship is not clear. However, at the conclusion of the workshop, scores on the post-workshop questionnaire rose an average of 5.8 points on a scale of one to 10, with 10 indicating the most favorable response (**Figure 2, Figure 3**, and **Figure 4**). These results suggest that, although the students were initially unenthusiastic about participating in the workshop or lacked an understanding of the graduate school process, they ultimately found it to be informative and worthwhile (**Figure 2A**, post-test). Prior to the workshop, students reported having little knowledge of the characteristics of graduate school (**Figure 2B**) and feeling unprepared for the application process (**Figure 2C**). Amidst their uncertainties, the students still did not think the workshop would be beneficial (**Figure 2D**), emphasizing the need for more “preparedness training”. This training refers to graduate school-specific information (e.g., the application process, interviews, mentoring) or rather efforts to convince students that graduate school is attainable with the proper preparation. It is possible that the students did not initially see the benefit of the workshop because they felt graduate school in general was beyond their grasp, thus making the workshop seem irrelevant to their goals, even if it did provide useful information. The students also did not expect the workshop to provide much help and clarity with the documents and experiences vital to the graduate school application process. Despite these low expectations, responses to the post-training questionnaire indicated that the students felt the information presented in the workshop empowered them to draft their personal statements (**Figure 3A**), seek letters of recommendation (**Figure 3B**), increase their confidence to apply to graduate school (**Figure 3C**), improve their GRE preparation, (**Figure 3D**) increase their networking skills (**Figure 3E**), and understand the value of research experiences, such as summer internships (**Figure 3F**). The post-training questionnaire results also indicated that the students felt the workshop was beneficial for their time management skills (**Figure 4A**), their ability to approach interview questions (**Figure 4B**), their understanding of the role of support teams (**Figure 4C**), their communication skills (**Figure 4D**), and their ability to identify and improve their learning styles (**Figure 4E**).

**Figure 2.**
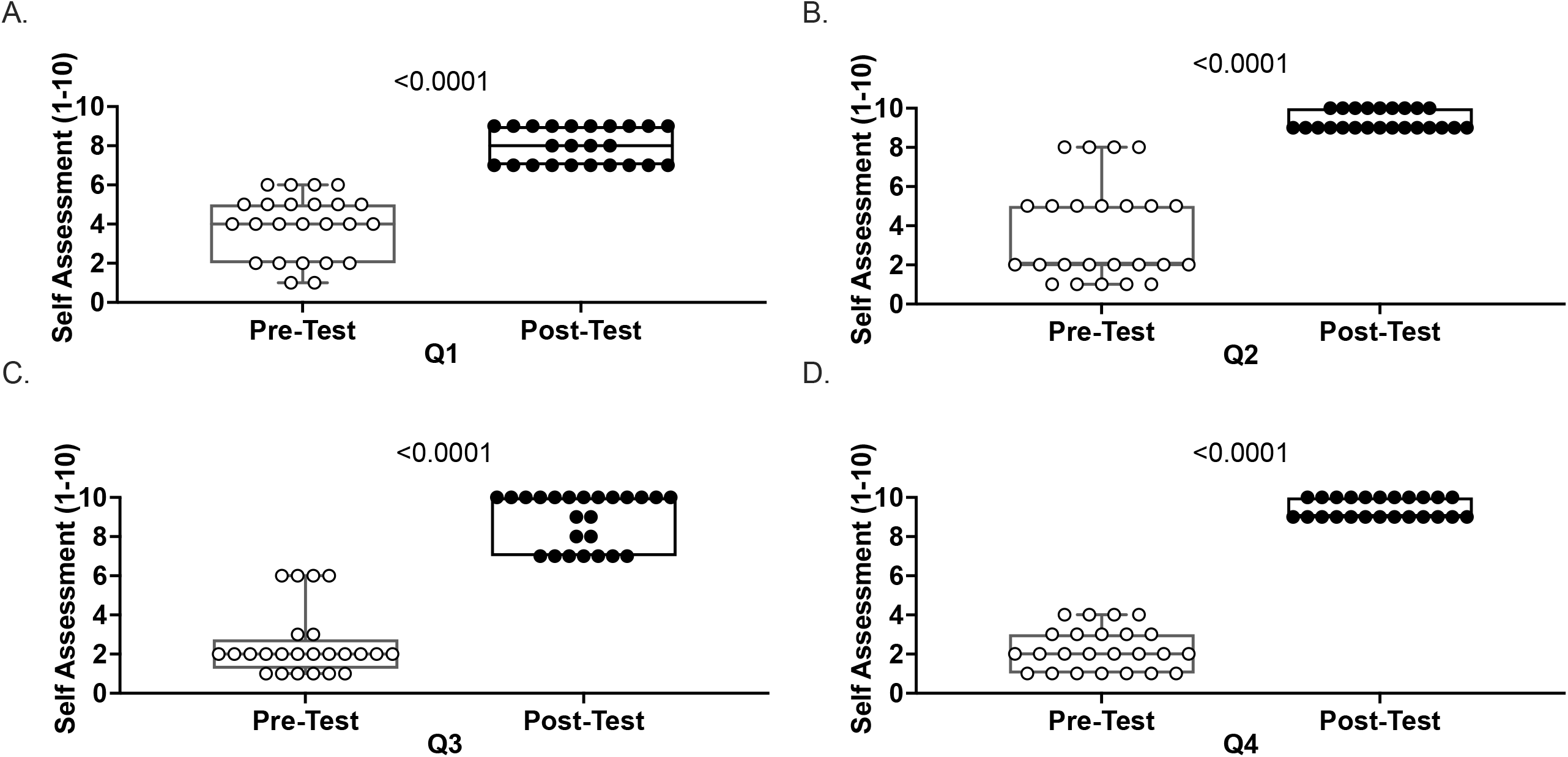
Results from pre- and post-workshop evaluations for questions 1–4 on their overall knowledge of graduate school.

**Figure 3.**
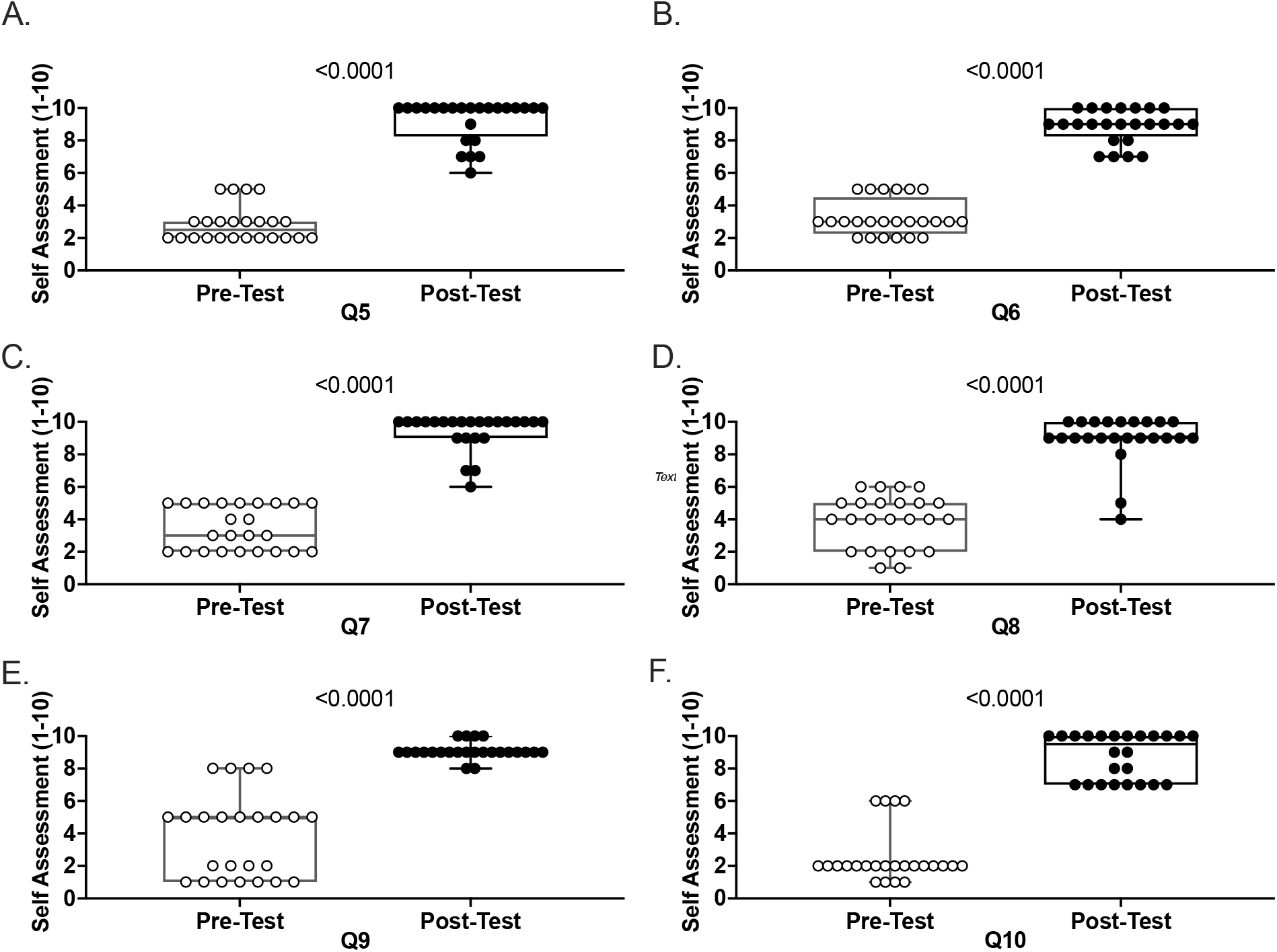
Results from pre- and post-workshop evaluations for questions 5–10 on how well the workshop prepared the participants for the graduate school application.

**Figure 4.**
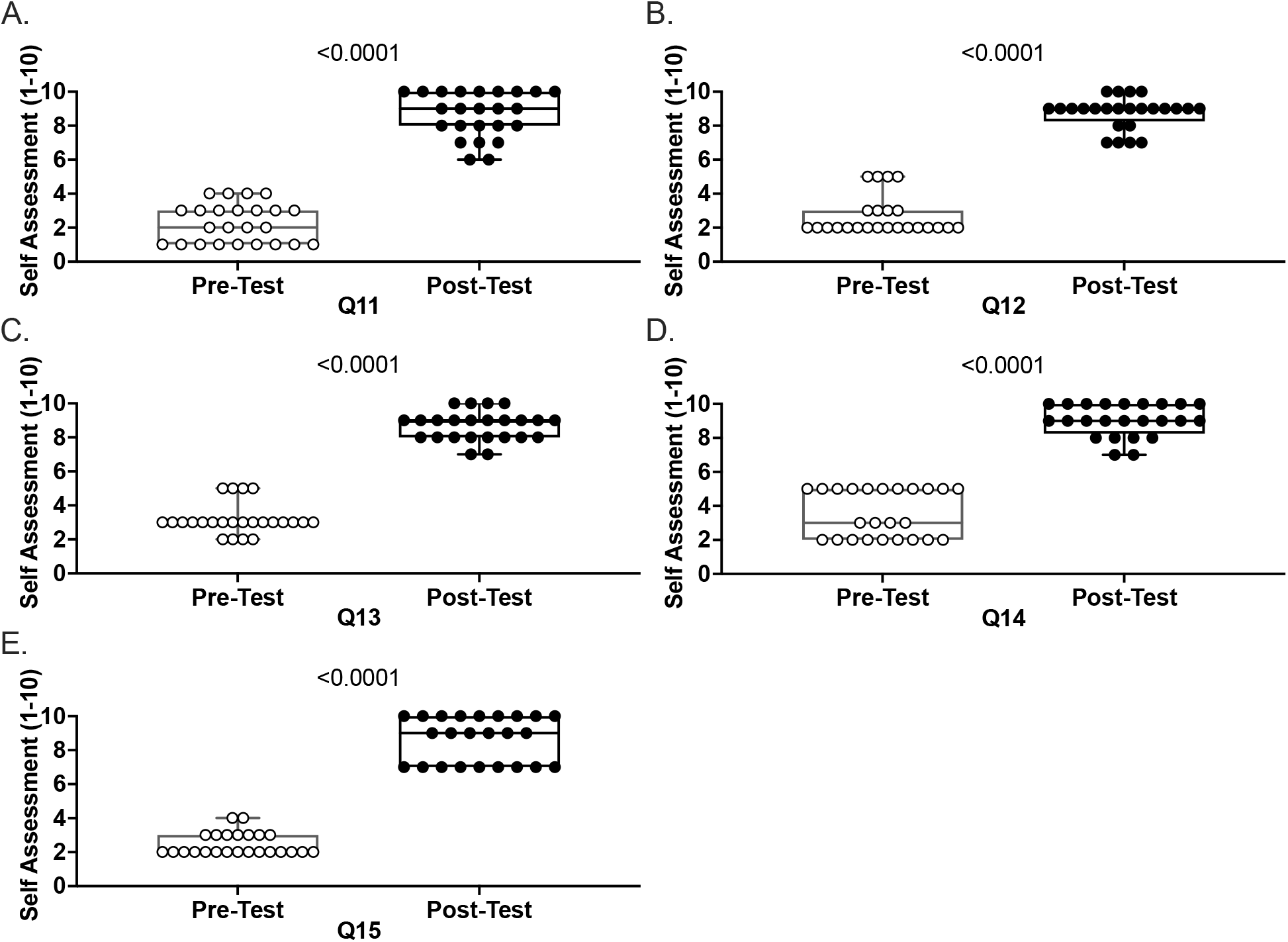
Results from pre- and post-workshop evaluations for questions 11–15 on overall preparedness for starting graduate school.

## Discussion

Collectively, the data from the questionnaires highlight the need for more career development opportunities, such as the workshop conducted in this paper. Although initial enthusiasm for this type of programming may be low, students can still benefit. Early exposure, mentoring, and incorporation of a champion’s mindset are all strategies that motivate and improve academic outcomes for URM students [14] [3, 4, 7-13, 15-20]. Though our sample size of surveyed students was small, our results suggest that many undergraduate URM students will also benefit from career development resources that aim to increase the influx and retention of URMs into the STEM pipeline, especially for those at minority-serving institutions with low retention rates. Success is shown to be more prevalent when students are empowered and equipped with the proper knowledge, tools, and strategies to overcome the overwhelming challenges of graduate school.

### Limitations

Due to the small number of participants, caution should be exercised in generalizing the results of this study. We also recognize that our study participants likely do not represent all STEM areas, as only those with a special interest in biology might have volunteered to participate. Moreover, participants may have changed their behavior after the workshop only because they felt obliged to fulfil expectations, rather than a true intent to pursue a career in STEM. Additional questions might have allowed a more comprehensive perspective of the students’ feedback.

## Availability of data and materials

A PowerPoint presentation of the workshop is available in English and Spanish formats. Survey data may be made available upon reasonable written request.

## Abbreviations

CV: Curriculum Vitae
GPA: Grade Point Average
GRE: Graduate Record Examinations
IDP: Individual Development Plan
MARC: Maximizing Access to Research Careers
MSI: Minority Serving Institution
NIH: National Institute of Health
NSB: National Science Board
NSF: National Science Foundation
QE: Qualifying Exam
RISE: Research Initiative for Scientific Enhancement
SROP: Summer Research Opportunity Program
REU: Research Experiences for Undergraduates
STEM: Science, Technology, Engineering, and Mathematics
URM: Underrepresented Minority

## Acknowledgements

We also would like to thank the 24 students who participated in our survey and will remain anonymous.

We would like to thank Heather Beasley for helping make the figure in BioRender.

## Funding

This work was supported by the UNCF/BMS EE Just Faculty Fund, Burroughs Wellcome Fund Award Career Awards at the Scientific Interface (CASI), NIH SRP subaward to #5R25HL106365-12 from the NIH PRIDE Program, BWF Ad Hoc Grant to D.M. and AHJ, and the Ford Foundation to A.H.J.; 1K99GM141449-01 MOSAIC grant to C.P.M. and NSF grant MCB #2011577I to S.A.M.

## Competing interests

Authors declare that they have no competing interests.

## Data and materials availability

All data are available in the main text or the supplementary materials.

## Ethics Declaration, Project Title

Promoting Engagement in science for underrepresented Ethnic and Racial minorities (P.E.E.R), 21-MortonD-HSR-SOM-01, Kaiser Foundation Research Institute FWA: FWA00002344

Ethics Approval and consent to participate, Yes

Consent for publication, Yes

Competing Interests, Not Applicable

## Notes

### Competing Interest Statement

The authors have declared no competing interest.

## References

1. Arbeit, C., Yamaner, M.I., National Center for Science and Engineering Statistics (NCSES), Trends for Graduate Student Enrollment and Postdoctoral Appointments in Science, Engineering, and Health Fields at U.S. Academic Institutions between 2017 and 2019. 2021: Alexandria, VA: National Science Foundation.

2. National Science Board, Vision 2030. 2020: Washington, D.C.

3. Hinton, A.O., Jr., et al., Patching the Leaks: Revitalizing and Reimagining the STEM Pipeline. Cell, 2020. 183(3): 568–575.

4. Stelter, R.L., J.B. Kupersmidt, and K.N. Stump, Establishing effective STEM mentoring relationships through mentor training. Ann N Y Acad Sci, 2021. 1483(1): 224–243.

5. Marshall, A., et al., Responding and navigating racialized microaggressions in STEM. Pathog Dis, 2021. 79(5).

6. Hinton, A.O., Jr., et al., Mentoring minority trainees: Minorities in academia face specific challenges that mentors should address to instill confidence. EMBO Rep, 2020. 21(10): p. e51269.

7. Shuler, H., et al., Intentional mentoring: maximizing the impact of underrepresented future scientists in the 21st century. Pathog Dis, 2021. 79(6).

8. Gordon, D.M., et al., Mentoring urban Black Middle-School Male Students: Implications for Academic Achievement. The Journal of Negro education, 2009. 78(3): 277–289.

9. Anderson, G.N., Dey, E.L., Gray, M., and Thomas, G., Mentors and proteges: The influence of faculty mentoring on undergraduates academic achievement, in Association for the Study of Higher Education. 1995: Orlando, Florida.

10. Thompson, L.A. and L. Kelly-Vance, The impact of mentoring on academic achievement of at-risk youth. Children and Youth Services Review, 2001. 23(3): 227–242.

11. DeFreitas, S.C., Bravo, Antonio, Jr, The Influence of Involvement with Faculty and Mentoring on the Self-Efficacy and Academic Achievement of African American and Latino College Students. Journal of the Scholarship of Teaching and Learning, 2012. 12(4): 1–11.

12. Honicke, T. and J. Broadbent, The influence of academic self-efficacy on academic performance: A systematic review. Educational Research Review, 2016. 17: 63–84.

13. Turner, E.A., M. Chandler, and R.W. Heffer, The Influence of Parenting Styles, Achievement Motivation, and Self-Efficacy on Academic Performance in College Students. Journal of College Student Development, 2009. 50(3): 337–346.

14. Strayhorn, T.L., Factors Influencing Black Males’ Preparation for College and Success in STEM Majors: A Mixed Methods Study. The Western Journal of Black Studies, 2015. 39(1).

15. Green, A. and D. Sanderson, The Roots of STEM Achievement: An Analysis of Persistence and Attainment in STEM Majors. The American Economist, 2018. 63(1): 79–93.

16. Kricorian, K., et al., Factors influencing participation of underrepresented students in STEM fields: matched mentors and mindsets. International Journal of STEM Education, 2020. 7(1): 16.

17. Dweck, C.S., Motivational processes affecting learning. American Psychologist, 1986. 41: 1040–1048.

18. Dweck, C.S., Mindset: The new psychology of success. 2006, New York, NY: Random House.

19. Rattan, A., et al., Leveraging Mindsets to Promote Academic Achievement: Policy Recommendations. Perspect Psychol Sci, 2015. 10(6): 721–726.

20. Hernandez, P.R., Schultz, P. W., Estrada, M., Woodcock, A., & Chance, R. C., Sustaining optimal motivation: A longitudinal analysis of interventions to broaden participation of underrepresented students in STEM. Journal of Educational Psychology, 2013. 105(1): 89–107.

