## Supplemental file for "Using a Champion-Oriented Mindset to Overcome the Challenges of Graduate School"

### The Trials and Tribulations of Graduate School:

#### How Do You Make an Impact?

### Outline

- Suggestion: Begin with a positive affirmation or oath promoting positivity
- Insert: Personal journey to graduate school
- How do you make the journey count ? Renewing your mindset to a Champion’s Mindset
- Structure of Graduate School
- Trials and Tribulations of Graduate School: Year One and Year Two
- What is necessary to get into graduate school?
- Suggestion: End with another affirming statement or oath

### Beginning Oath or Affirmation

*Insert a positive affirmation or oath promoting positivity*

### How Do I Get Into Graduate School?
### Step 1: A Champion's Mindset!

Insert Inspirational Quote Here

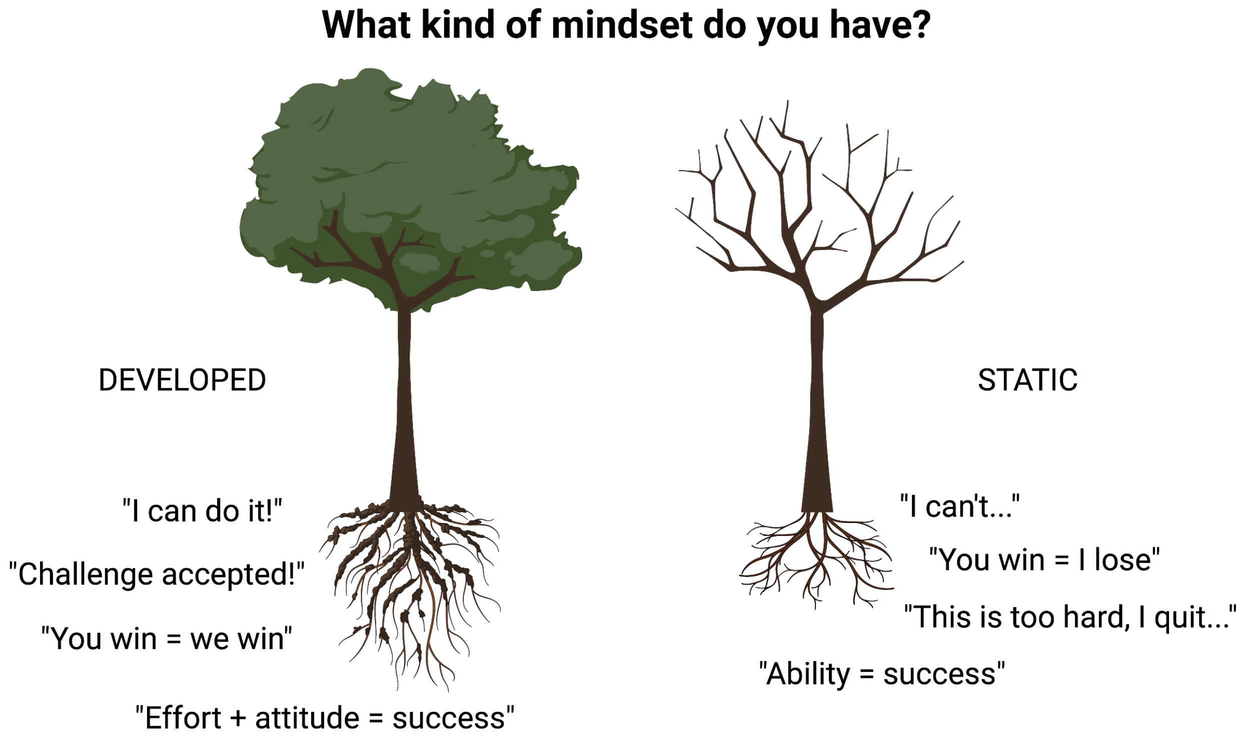


I want each of you to be able to access the power inside of you to achieve a pleasant life and the indomitable success we all look for in our educational plan.

### How Do I Get Into Graduate School?
### Step 1: A Champion's Mindset!

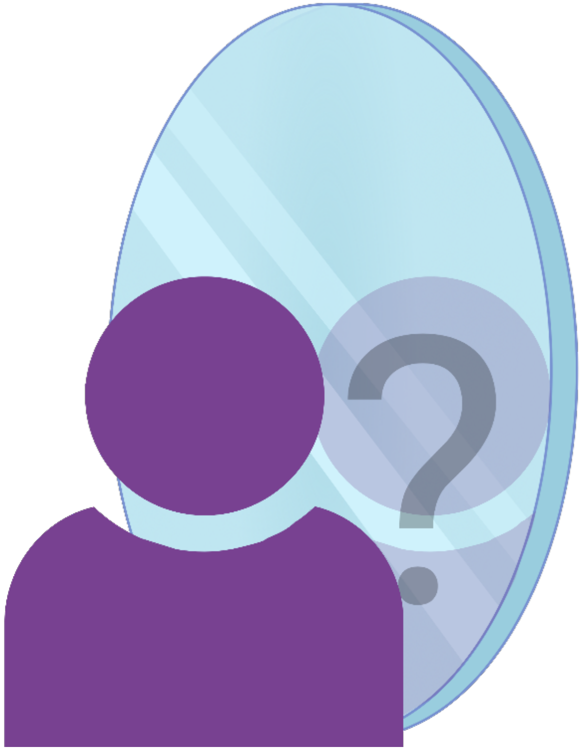


First thing to do is find out who you are and what you want in life!

- Image
  - If you do not have role models or mentors, you cannot set an image for yourself.
  - You will have a hard time telling your story.
  - We all need a guide!
- Identity
  - How do you develop a strong identity?
  - Enjoy the journey!

Insert Inspirational Quote Here

### How Do I Get Into Graduate School?
### Step 1: A Champion's Mindset!

Consider inserting images of your mentors or others who have inspired you

First thing to do is find out who you are and what you want in life!

***Consider inserting an inspirational quote***

Image

- If you do not have role models or mentors, you cannot set an image for yourself.
- You will have a hard time telling your story.
- We all need a guide!

Identity

- How do you develop a strong identity?
- Enjoy the journey!

### How do I get into Graduate School?
### Step 1: A Champion's Mindset!

Consider inserting images of your role models

First thing to do is find out who you are and what you want in life!

*Consider inserting an inspirational quote*

- Image
  - If you do not have role models or mentors, you cannot set an image for yourself.
  - You will have a hard time telling your story.
  - We all need a guide!
- Identity
  - How do you develop a strong identity?
  - Enjoy the journey!

### How Do I Get Into Graduate School?
### Step 1: A Champion's Mindset!

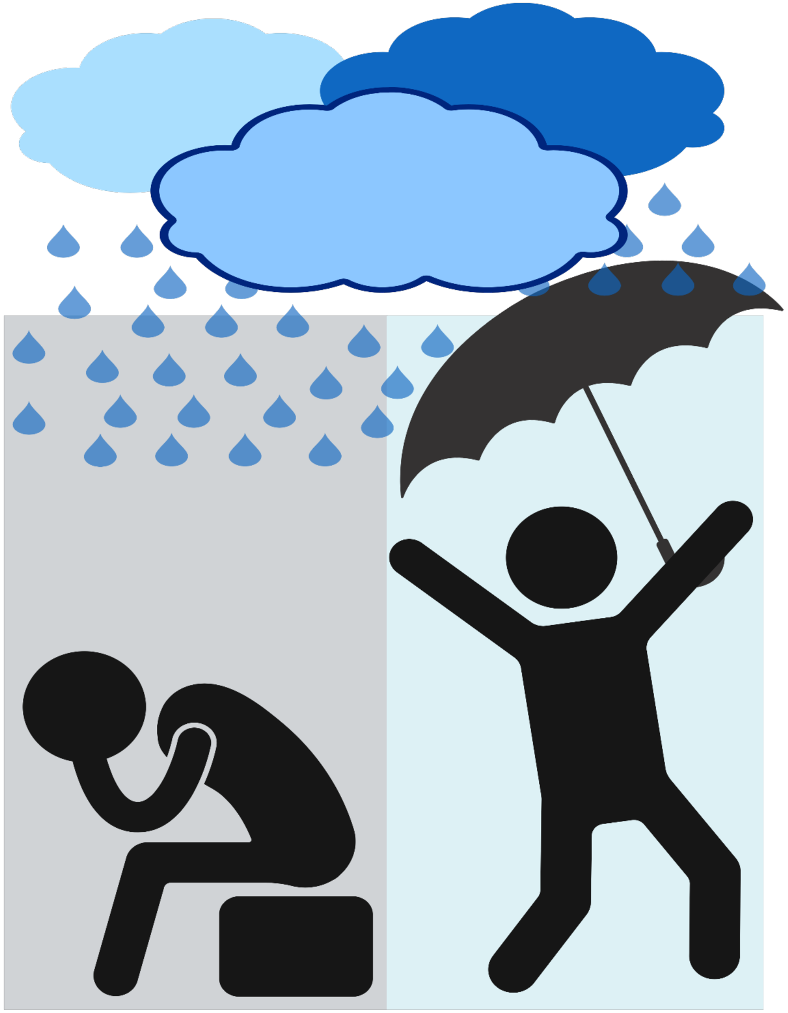

- Being Proactive
  - When adversity comes, change how you deal with it!
  - What happens if you do not know how to deal with it ?
- Power of your words.
  - Difference between saying, "I can" and "I will" versus, "I will try to do this."

**Insert inspirational quote here**

### How Do I Get Into Graduate School?
### Step 1: A Champion’s Mindset!

Consider inserting images of overcoming adversity.

- Being Proactive
  - When adversity comes, change how you deal with it!
  - What happens if you do not know how to deal with it ?
- Power of your words.
  - Difference between saying, “I can” and “I will” versus, “I will try to do this.”

**Insert inspirational quote here**

### Why Do You Need A Champion's Mindset?

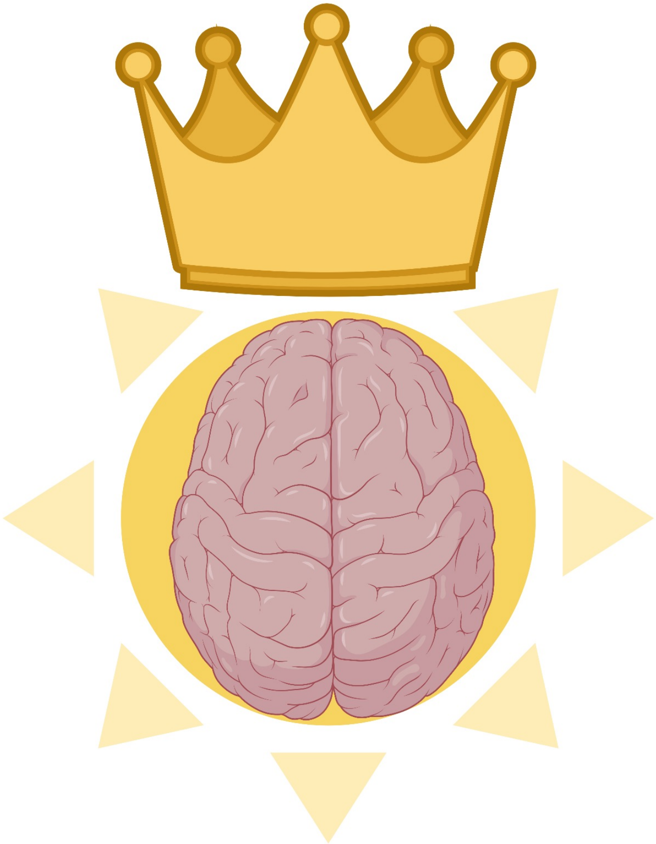

- Resolve
- Perseverance
- Access to Great Verbal and Non-verbal communication
- To unlock your potential

**Insert inspirational quote here**

### Structure of Graduate School

- **Summer Before-** Have as much fun as possible. I also suggest one month before going to grad school try to acquire access to a syllabus, old notes, and old test banks to help with your studying.
- **1st year** –Classes or Modules and Rotations
- **2nd year** –Classes, Modules, Qualifying Exam
  - (depending on school, you could have two qualifying exams), Picking a laboratory or the lab will pick you.
- **3rd year** –Classes and Research in desired laboratory, Qualifying Exam, Career Development should start here (IDP), Conferences on Career Development, Grant Writing, and Graduate Grant.

Consider adding personal images of fun events

### Structure of Graduate School

- **4th year** – Research, Career Development, Thinking about your next steps, Internship for summer, if interested in a non-academic field, writing up one first author (at least).

- **5th year**– Thinking about your next program, setting up your career, final career development goals, finding your niche and asking others to collaborate and acquire extra papers.

- **6th year** and Beyond– Getting out !!!

### Year One Trial and Tribulations- Classes

**Deficiencies show up**

- Classes in undergraduate studies that were difficult.
- Classes in undergraduate studies that you avoided because of the rigor.
- Not being able to work together in a group or small study group.

**Letting go of who you were to become who you want to be in the future.**

- From Undergrad, if you were the big fish in a small pond or a big fish in a large pond, realize that graduate school is a different beast.
- Understand that it is okay to be humble, as it is the best thing needed to be successful, because people will perceive you as teachable.

### Year One Trial and Tribulations- Stress

#### Stress and Time Management

- Learning more about your personality.
- Understanding the daily plan and how your week will go is important ( allow yourself time to plan in advance).
- Stress will occur and it is okay to talk and hangout with friends.
- Stress also can become overwhelming so getting counseling can help you.

#### Classes are Priority, not Lab the first year, but you still need to make substantial progress

- A mentor will sometimes ask you to collect data and put aside your study time.
- Important Advice: you have to find the balance of what can work for your rotation because you have to please your PI to get into the laboratory, but you do not have time to sacrifice your classes. Academic Probation is worse!

### Year One: Trial and Tribulations- Relationships

#### Friend Dynamics Change

- Through out graduate school you have constant ebb and flow with relationships.
- Be aware when a relationship is not working for you, do not be afraid to change it. Your health in graduate school is key to your success.
- Take the time to find friends to release stress.
- Until you make friends, try to focus on hanging out with your cohort, as these are the people you will study with for the semester (Early Networking).

#### Family Relationship dynamics change

- This is not a four year degree, so when your family asks for a graduation date, tell them I will let you know around year 4 or 5. This type of pressure to graduate can cause for additional stress.
- Family deaths can impact how you manage your time and what your perception of reality is like at the moment.
- Major relationships can also influence your performance for the better or for the worse.
- Family vacation is important as well. Stay connected to your family as mental health is important.

### Year Two: Trial and Tribulations- Qualifying Exam

**Understanding your Capacity to Learn**

- How do you learn?
- Are you sure you understand what you are studying ?
  - Who
  - What
  - When
  - Where
  - Why
  - How

**Determining your focus**

- What is more important ?
  - Your goal or an individual achievement?
  - Your paper or weekend off ?
  - Studying or really practicing for the qual?

### What Is Necessary For My Graduate Acceptance ?

- Individual Development Plan (IDP)
- Letters of Recommendation
- Grade point average (GPA)
- Optional: Graduate Record Examination (GRE)
- Summer Research Programs
- Masters or Post-bac Programs
- Conferences
- Undergraduate Productivity- Honors & Awards, Shadowing, Volunteering, Participating in on Campus Organizations, and Showing Leadership Potential

### IDPs For The Undergraduate Level

- **What is an IDP ?**
  - Individual Development Plans (IDPs)
- **Why do we need an IDP?**
  - Setting goals can help you be more intentional about the experiences you have in your undergraduate or graduate education and training, and can provide key steps in the right direction for you. The best goals are specific, measurable, achievable, relevant, helps facilitate clear communication, and timely.
- **What can an IDP do for me ?**
  - To Assess your current skills and strengths
  - To create a plan for developing and enhancing skills to help you meet your academic and professional goals
  - To communicate with your mentors about your evolving goals and related skills
- **Who needs an IDP ?**
  - We all can use one!

### Individual Development Plan (IDP) For Undergraduate Level and Transitioning To A Graduate Level (IDP)

**Undergraduate Level**

- Includes an Assessment of your Personality
  - Big Five Personality Test

    (Five Factor Model-Openness, Neuroticism (feeling)j, Agreeableness, Extraversion, Conscientiousness)
  - Myers-Briggs Personality Types
    - ENFJ ( MLK, Obama, Oprah, David, Peyton, and Sampras)
    - Important Values : Home/Family, Health, Friendship, Financial Security, and Learning
    - Important traits: Imbue others, enthusiastic, want others to leave up to their potential.

**Undergraduate Level**

- Includes an Assessment of your Love Language
  - Words of Affirmation
  - Acts of Service
  - Receiving Gifts
  - Quality Time
  - Physical Touch

### Individual Development Plan (IDP) For Undergraduate Level and Transitioning To A Graduate Level (IDP)

#### Undergraduate Level

- Includes an Assessment of your Learning Styles

1. Visual -See
2. Auditory – Sound & Music
3. Verbal –Speech
4. Physical – Hands on
5. Logical –Reasoning and logic
6. Social-learn in group or with other people.
7. Solitary-work alone and use self-study

### Individual Development Plan (IDP) For Undergraduate Level and Transitioning To A Graduate Level (IDP)

**Undergraduate Level**

- Career Development
  - Internships (Amgen, DAAD, CIC, and Leadership Alliance)
  - Join Research gate & LinkedIn
  - Join Honors Society & Campus Organization
  - Join Making Business Cards
  - Networking for opportunities
    - Conferences
      - EB
      - Posters on the Hill
      - International Conference of Undergraduate research
      - National Conference on undergraduate research

**Undergraduate Level**

- Teaching and Tutoring
  - Teacher Assistant
  - Tutoring
  - Mentor students
- Short Term Aspirations
  - Finish undergraduate school
  - Win Award
  - Present a poster
- Long Term Aspirations
  - Ex. Graduate, Medical, or Professional School
  - Ex. Job directly afterward

### Individual Development Plan (IDP) For Undergraduate Level and Transitioning To A Graduate Level (IDP)

#### **Weaknesses and Strengths**

#### What Are My Strengths?

- Insert a list of your personal strengths during this transition time

### Individual Development Plan (IDP) For Undergraduate Level and Transitioning To A Graduate Level (IDP)

**Weaknesses and Strengths**

- What do I need to improve?
  - Include a list of your personal weaknesses during this transition period

### Individual Development Plan (IDP) For Undergraduate Level and Transitioning To A Graduate Level (IDP)

#### Undergraduate Level

**Research Goals:**

- To develop my laboratory skills
- To help with grant writing
- To work as an effective team in the laboratory with others.
- To read the literature and understand the global picture in my research
- To acquire a position on a paper

#### Undergraduate Level

**Awards and Honors:**

**Awards:**

- Research Award for undergraduate research
- Leadership awards
- Honors thesis award
- Dean's and Chancellor's List

**Fellowships:**

What fellowships should I write as a graduate student ?

### Individual Development Plan (IDP) For Undergraduate Level and Transitioning To A Graduate Level (IDP)

**National Fellowships**

- NIH F-30 (MD/Ph.D)
- NIH F-31 (Ph.D)
- NSF
- DOD NSDEG
- Mustard Seed Fellowship
- ADA
- AHA
- Department of Energy computational sciences fellowship
- Javits Fellowship

### Individual Development Plan (IDP) For Undergraduate Level and Transitioning To A Graduate Level (IDP)

#### National Diversity Fellowships

- Ford Foundation
- ADA Minority
- NIH supplement to an RO1
- F-31 Diversity
- UNCF MERCK
- F-31 Diversity
- Burroughs Wellcome Fund
- APS Porter Physiology Fellowship
- Harriett G. Jenkins Pre-doctoral Fellowship
- HHMI Gilliam Fellow
- Hertz Fellowship

### Individual Development Plan (IDP) For Undergraduate Level and Transitioning To A Graduate Level (IDP)

#### International Fellowships

- “Paul and Daisy Soros Fellowship – Just received permanent resident status or a green card
- HHMI fellowship- discontinued and not sure if it will continue
- AHA- accepts international applicants
- American Association of University Women International Fellowships
- Fulbright Program for Non-US Students
- Oxford and Rhodes Scholarships
- Gates Cambridge Scholars Program –Attend UK
- International Peace Scholarship (Women) – US and Canada

### GPA And Picking A Major That Works For You

- What is an acceptable GPA for graduate school?
  - 3.0 and greater
  - Semester Improvements
- Does it matter what University your GPA comes from and does your Science and Math GPA matter?
  - Yes
- How can I ameliorate my GPA ?

Special cases for getting into graduate school with out a 3.0.

- post-bac or Master's Program to augment performance
- Retake a course or use GRE subject test
- Show steady growth after semester that did not go well.

### GRE Scores

- Why should I take a GRE subject test?
  - Can demonstrate your mastery of a particular field of study
  - Can help you stand out from other applicants by emphasizing your knowledge and skill level in a specific area

- What is an acceptable GRE for graduate school?
  - Generally 75 percentile is acceptable at most schools and an average of 3.5 to 4 on writing.
  - Desired 90 percentile ( for all schools)
  - Can I still be competitive with a GRE score in the 50 percentile?
    - yes

### GRE Scores

- How do I study when I do not have the money to take a GRE course ?
  - Buy a study book
  - Make your own course!!!
  - Seek published sources online

Insert example questions here

Insert example questions here

### GRE Scores

Insert example problems here

### Personal Statements

Introduction: What makes a good introduction ?

Fourteen years ago, I was relaxing on my parents' couch watching TV, when my mom came home with an interesting gift for me from one of her coworkers. It was the book entitled *The Hot Zone* by Richard Preston. At the age of 15, I was typically used to reading cheesy mysteries involving overly dramatized characters and cliché endings. Something about this book intrigued me though. I began reading it from cover to end and instantly fell in love. This bio-thriller immediately opened my eyes to a world of science that was so foreign to me. Seeing as how my parents were completely ambivalent toward the sciences and never encouraged any learning of the discipline, my exposure was limited to that of boring and insufficient hour-long school lectures. An indomitable passion for the sciences was ignited in me following the completion of this novel and has yet to cease all these many years.

Word Choice
Interesting Hook
Describing how Science Impacted her in a clever way

Published with permission of XXX

### Personal Statements

How should I address my deficiencies?

I have had the pleasure of attaining an undergraduate degree in Biology from the University of Houston-Downtown, while raising two beautiful girls. Though working towards pedagogic excellence in the natural sciences and trying to imbue strong values in my children has not been an easy assignment, my extensive life experience has instilled in me a unique sense of maturity and strong work ethic. I truly had to embody these two characteristics to endure the lengthy and challenging academic road that awaited me. Despite having to retake a few courses and delaying my graduation date by three years, I was able to finish in the top 10 percent of my class and was inducted into Scholars' Academy, the Honors Society for the Sciences.

She addressed that it took her three additional years to acquire her undergraduate degree.

She indirectly addressed the achievement gap that some professors in science think that women are not as capable.

### Personal Statements

How should I address my deficiencies?

Academy, the Honors Society for the Sciences. I also faced an additional challenge when confronted with the GRE. I was stricken with extreme test anxiety and my performance was not an accurate reflection of my true abilities. In an effort to surmount these hurdles, I applied and was accepted to the SMART PREP post baccalaureate program at Baylor College of Medicine where I was presented with a course in molecular and cell biology, a course in graduate presentation skills, a chance to work in a well known lab along side experienced post docs, and a math tutor to strengthen my math skills. At Baylor, I learned how to think critically, read and analyze scientific data, and master effective presentation skills. By persevering through difficult challenges and taking the right steps to improve myself as a scientist, I have demonstrated that minor academic mistakes and a score on a standardized test do not by themselves define my potential success as a student. I have full confidence that I will be able to complete a Ph.D. program using both my undergraduate research and post baccalaureate experiences as a strong foundation.

Addresses her issues with the GRE
Addresses by getting a math tutor

Addresses her lack of exposure to research by going to the SMART PREP program
(Important to note, during her post-bac she published a paper)

### Personal Statements

How do I show that I am unique and can improve the PhD program?

In addition, having to balance multiple responsibilities, as a mother, has also strengthened my time-management skills and taught me that I must be patient and cogent in my endeavors to ensure good results. These attributes have followed me throughout my academic career and serve to fortify my accord in the lab. As a researcher, my domestic life as well as my experience in the lab has served to shape my understanding of the empirical methods involved in conducting research and has strengthened my ability to perform various tasks in a laboratory setting. At the bench, I am steady and deliberate

with my handling techniques. Because of this, I acquire consistently strong results. My love for learning has allowed me to pick up on new skills relatively quickly and I am then able to put them into action on critical studies. Through my training in various techniques and good mentorship I have become a self-directed learner and I have acquired knowledge that transcends classroom study.

She decided to demonstrate how being a mother allows her to acquire strong and consistent results

### Letter of Recommendation

How do I know that my professor is
writing a strong letter of recommendation for me ?

- Classes the student has taken with the recommender.
- Experiences you have shared.
- Transcripts.
- Resume/CV.
- Research experience and internships.
- Awards and achievements.
- Academic/career goals.
- Relevant professional experience.

### Letter of Recommendation

How do I know that my professor is
writing a strong letter of recommendation for me ?

#### Adjectives to Avoid :

caring
compassionate
hard-working
conscientious
dependable
diligent
dedicated
tactful
interpersonal
warm
helpful

#### Adjectives to Include

successful
excellent
accomplished
outstanding
skilled
knowledgeable
insightful
resourceful
confident
ambitious
independent
intellectual

### Letter of Recommendation

How do I know that my professor is
writing a strong letter of recommendation for me ?

**Language is key**

**Men and Women Letters are often different.**

**We have to avoid gender bias when writing letters.**

Trix, F & Psenka, C. Exploring the color of glass: Letters of recommendation for female and male medical faculty. Discourse & Society, 2003; and Madera, JM, Hebl, MR, & Martin, RC. Gender and letters of Recommendation for Academia: Agentic and Communal Differences. Journal of Applied Psychology, 2009.

### Letter of Recommendation

How do I know that my professor is
writing a strong letter of recommendation for me ?

**Language is key – Keep It Professional**

**Letters of Rec for women are more likely to mention personal life .**

**Personal should be irrelevant**

**Formal titles an surnames should be for men, women, or gender neutral pronouns.**

Trix, F & Psenka, C. Exploring the color of glass: Letters of recommendation for female and male medical faculty. Discourse & Society, 2003; and Madera, JM, Hebl, MR, & Martin, RC. Gender and letters of Recommendation for Academia: Agentic and Communal Differences. Journal of Applied Psychology, 2009.

### Letter of Recommendation

How do I know that my professor is
writing a strong letter of recommendation for me ?

**Letters of Rec for men are more likely to mention publications**

**Letter of Rec for men are more likely to state multiple references to research**

**Make Sure you put these critical accomplishments in every letter.**

Trix, F & Psenka, C. Exploring the color of glass: Letters of recommendation for female and male medical faculty. Discourse & Society, 2003; and Madera, JM, Hebl, MR, & Martin, RC. Gender and letters of Recommendation for Academia: Agentic and Communal Differences. Journal of Applied Psychology, 2009.

### Letter of Recommendation

How do I know that my professor is
writing a strong letter of recommendation for me ?

**Letters of rec for men are more likely to emphasize accomplishments**

**Make sure women's letters of rec have effort but also ability.**

**Do not associate stereotypes of women in the letter.**

**Do not raise doubts in letters. Women are more likely for this to happen.**

Trix, F & Psenka, C. Exploring the color of glass: Letters of recommendation for female and male medical faculty. Discourse & Society, 2003; and Madera, JM, Hebl, MR, & Martin, RC. Gender and letters of Recommendation for Academia: Agentic and Communal Differences. Journal of Applied Psychology, 2009.

### Interview Questions

### Interview Questions

Remember, your goal is to convey your curiosity, enthusiasm and competence, but also glean information **YOU** need to conclude if this graduate program is a good fit for you.

**You should know what they are not allowed to ask you:**

Questions that focus on a person's race, creed, religion, gender, sexual orientation, position on a political matter, marital status, one’s age, physical/mental disability or learning disability, remember it is illegal.

### Interview Questions

**How to prepare:**

- Review research of all faculty you are interviewing with, know who the Dean is for the graduate program you are applying to, know who the Diversity Dean or Director is for that program, and lastly, make sure you aware of some of the star students in your program and the graduate college. Additionally, make sure you can address all questions related to deficiencies in your application.
- I want to stress that it is important to obtain adequate practice before you go on your interviews. Therefore, Practice, practice, practice; mock interviews!!!
- After each mock interview, take the time to review recommendations and "rehearse potential answers" that you stumbled on in the mock.
- Prepare at least 2 questions for EACH person you will interview with related to the graduate program, the interviewers' research, and ask about their experience with the program.
- List questions about the specific program that are NOT found on the website, thus you need to prepare by going to the graduate website and knowing the program inside and out.

### Interview Questions

**Personal Characteristics:**

1) Tell me a little about yourself.
2) What are your biggest strengths?
3) What are some of your weaknesses?
4) What do you believe your greatest challenge will be if accepted into this program?
5) Describe your greatest scientific accomplishment so far.
6) What do you do in your spare time for fun?
7) Why should we take you and not someone else?
8) How would your professors describe you?

### Interview Questions

**Academic experiences and skills:**

1) In college, what course did you enjoy most? Least & why?
2) Describe your most current research project.
3) Why did you choose to apply to our program?
4) What do you know about our program?
5) What other schools are you considering?
6) In what ways have your previous experiences prepared you for graduate study in our program?
7) Tell me about your experience in the field? What was challenging about gaining competence in your research field of interest? What were your contributions so far to the field and what impact do you think you will make while in graduate school?

### Interview Questions

**Personal Characteristics:**

1) Tell me a little about yourself.
2) What are your biggest strengths?
3) What are some of your weaknesses?
4) What do you believe your greatest challenge will be if accepted into this program?
5) Describe your greatest scientific accomplishment so far.
6) What do you do in your spare time for fun?
7) Why should we take you and not someone else?
8) How would your professors describe you?

**Academic experiences and skills:**

1) In college, what course did you enjoy most? Least & why?
2) Describe your most current research project.
3) Why did you choose to apply to our program?
4) What do you know about our program?
5) What other schools are you considering?
6) In what ways have your previous experiences prepared you for graduate study in our program?
7) Tell me about your experience in the field? What was challenging about gaining competence in your research field of interest? What were your contributions so far to the field and what impact do you think you will make while in graduate school?

### Interview Questions

**Problem solving and leadership skills:**

1) Can you tell me about a problem you have had to overcome and the steps you took to resolve it?
2) How well do you handle stress? Can you provide an example?
3) How well do you manage your time? Can you provide an example?
4) How do you overcome micro-aggressions with individuals that are not used to interacting with people of color?

### Interview Questions

**Goals:**

1) Why do you need a Ph.D. to accomplish your career goals?
2) What drove your interest in science?
3) What excites you about a career in science?
4) If you didn't get accepted by any graduate schools, what would be your plans?
5) What are your career goals in 5 years and in 10 years

### Interview Questions

**How do you respond to an inappropriate question?**

1. You can say, “I do not feel comfortable answer this question.”

2. You can also state that you understand the question and though it may be illegal, you are okay with answering the question. Yet when you do this, you open yourself up to a certain line of questioning that could end up making you feel very uncomfortable. Try your best to avoid this line of questions!

### Questions??

Questions ???
